# β-Hydroxybutyrate maintains energetically demanding neural functions during glucose deprivation

**DOI:** 10.64898/2026.06.07.730699

**Authors:** Rachel Ricks, Dallin S. Nevers, Tyler Poulos, T. Luke Shafer, Clayton Dunford, Paul R. Reynolds, Benjamin T. Bikman, R. Ryley Parrish

## Abstract

Ketone bodies are a major source of cerebral energy during fasting and the ketogenic diet, but whether they can independently sustain brain tissue metabolism when glucose is absent remains uncertain. This question is difficult to resolve *in vivo* because circulating glucose is maintained, even during starvation, through endogenous production. We therefore used an *ex vivo* brain preparation to examine the metabolic capacity of tissue supplied with β-hydroxybutyrate (BHB) as the only exogenous fuel, isolating brain tissue from the primary endogenous glucose sources. Mitochondrial function was monitored following prolonged exposure to glucose-free, BHB-rich artificial cerebrospinal fluid, and tissue resilience was tested by inducing spreading depolarization, a severe energetic challenge that requires rapid restoration of ionic and metabolic homeostasis. Following acute reliance on BHB, mitochondria appeared to dynamically regulate electron transfer system function, utilizing lower O_2_ flux, while maintaining sufficient energetic reserves to preserve tissue ability to generate and recover from repeated spreading depolarizations. These findings demonstrate that BHB can independently maintain essential metabolic and functional properties of brain tissue in the absence of exogenous glucose. The results broaden our understanding of cerebral fuel flexibility and provide additional support for the use of ketogenic strategies in neurological disorders where tissue excitability, energy metabolism, or glucose availability may be altered.

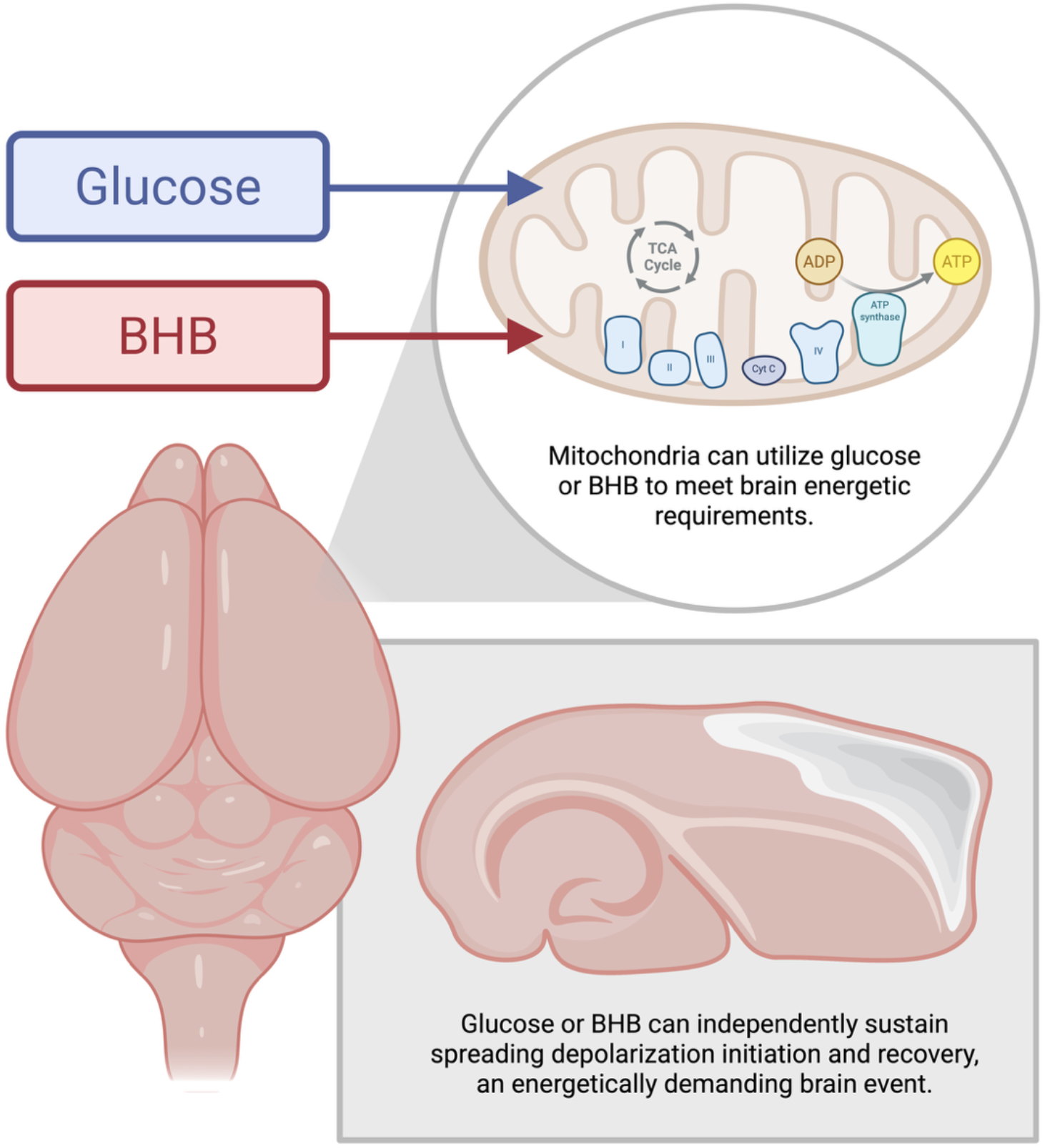

**Cerebral fuel flexibility: glucose and ketones:** β-Hydroxybutyrate (BHB) supports continual mitochondrial oxygen flux, along with induction of and recovery from repeated spreading depolarizations in mouse brain tissue in the absence of glucose. These findings demonstrate that ketone-supported metabolism can sustain energetically demanding neural function through a distinct bioenergetic strategy. Created in BioRender. Parrish, R. (2026) https://BioRender.com/k92j91u.

## Introduction

The mammalian brain is classically viewed as a glucose-dependent organ, having a high energetic demand and limited energy stores. Therefore, under normal physiological conditions, it is believed that glucose is the predominant substrate supporting cerebral ATP production (Dienel 2019; Mergenthaler et al. 2013). However, during prolonged fasting, ketogenic diet adherence, or other states of reduced carbohydrate availability, circulating ketone bodies increase and become an important alternative fuel for the brain, a state known as ketosis (Bolaños and Magistretti 2025; Chowdhury et al. 2014; García-Rodríguez and Giménez-Cassina 2021). Among these alternative fuel sources, β-hydroxybutyrate (BHB) is the major circulating ketone body and can be transported across the blood–brain barrier and oxidized by neural cells to support mitochondrial energy production (Achanta and Rae 2017; Cahill and Veech 2003).

Although ketone bodies are widely recognized as glucose-sparing substrates, it remains unclear whether BHB alone can sustain brain tissue energetics in the near-complete absence of glucose. Defining how brain tissue responds to true glucose deprivation can aid in understanding therapeutic states of ketosis, particularly in conditions where it is widely used, such as drug-resistant epilepsy and GLUT1 deficiency syndrome (Bough and Rho 2007; Daley et al. 2026; Qiao et al. 2024; Williams et al. 2024). Creating states of ketosis, either by endogenous or exogenous means, is also increasingly being considered as a therapy for many other neurological conditions (Jensen et al. 2020; Luong et al. 2025). Therefore, direct experimental tests of brain function and viability under glucose-free, BHB-supported conditions can help to clearly define the limits of cerebral metabolic flexibility. In intact mammals, glucose can still be produced during prolonged starvation through gluconeogenesis, making it difficult to isolate the brain’s dependence on glucose versus alternative fuels *in vivo* (Rui 2014). However, this question can be answered using *ex vivo* brain preparations, where brain tissue can be isolated from almost all sources of gluconeogenesis.

In this study, we quantified brain tissue mitochondrial respiration when BHB is provided as the sole exogenous energetic substrate, along with cerebral ability to undergo and recover from arguably the most energetically demanding event that occurs in the brain, namely spreading depolarization (SD) (Lindquist 2024). Our data demonstrate that mitochondria acutely utilizing BHB employ lower O_2_ flux across respiratory states compared to glucose. In addition, we found that brain tissue can still generate multiple SDs in the absence of glucose, challenging the idea that glucose is necessary for brain metabolic function. These findings provide direct evidence for the capacity of BHB to serve as a standalone cerebral fuel and have implications for understanding ketogenic neuroprotection, metabolic resilience, and therapeutic strategies targeting brain energy metabolism.

## Materials and Methods

### Animal Care and Ethical Approval

All animal procedures were approved by the Institutional Animal Care Committee (IACUC) and conducted in accordance with institutional guidelines. Experiments were performed on male and female C57BL/6 mice aged 3–9 weeks. Animals were housed in individually ventilated cages under a 12-hr light/12-hr dark cycle with *ad libitum* access to food and water.

### Brain Slice Preparation

Mice were anesthetized with isoflurane and decapitated. Brains were immediately removed and immersed in ice-cold carbogenated (95% O_2_/5% CO_2_) cutting solution containing (in mM): 126 NaCl, 26 NaHCO_3_, 3.5 KCl, 3 MgCl_2_, 1.26 NaH_2_PO_4_, and 10 glucose. Horizontal brain slices (400 μm) were prepared using a VT1200 vibratome (Leica Microsystem) and further dissected into neocortical sections as previously described (Withers et al. 2025). Brain slices were then quickly transferred to a continuously carbogenated artificial cerebrospinal fluid (aCSF) bath containing (in mM): 126 NaCl, 26 NaHCO_3_, 3.5 KCl, 2 CaCl_2_, 1 MgCl_2_, 1.26 NaH_2_PO_4_, and 10 glucose. The prepared neocortical brain slices were then randomly assigned to one of the following solutions for experimentation where appropriate: aCSF containing either 10 mM glucose, 10 mM racemic β-hydroxybutyrate (BHB), or 10 mM sucrose. Experimental solutions containing BHB or sucrose were made by replacing glucose with equimolar sodium β-hydroxybutyrate or sucrose, while ensuring consistent osmolarity and pH between treatment solutions.

Brain slices used for high-resolution respirometry were maintained in a submersion chamber at 37°C for 4 hours prior to experimentation. Brain slices used for SD experiments were maintained in an interface holding chamber at room temperature for 3–8 hours in their respective solutions prior to experimentation.

### High-Resolution Respirometry

Following the 4-hour incubation in one of the three prepared aCSF solutions, brain slices were weighed using an analytical balance (Mettler Toledo AL104). Slices were then transferred to 1.7 mL microcentrifuge tubes containing ice-cold MiR05 respiration medium containing (in mM): 110 sucrose, 60 lactobionic acid, 20 taurine, 20 HEPES, 10 KH_2_PO_4_, 3 MgCl_2_, 0.5 EGTA, supplemented with 1 g/L BSA. Tissue was mechanically homogenized by repeated aspiration with a pipette. The resulting homogenate was transferred to an Oroboros O2k high-resolution respirometer (Oroboros, Innsbruck, Austria) containing MiR05 maintained at 37°C for evaluating mitochondrial fitness.

Following chamber loading, dissolved O_2_ concentration was elevated to 400 μM to establish baseline O_2_ availability. Mitochondrial respiration was then assessed using a substrate-uncoupler-inhibitor titration (SUIT) protocol. Each successive titration was performed after a stable O_2_ flux plateau was recorded.

Non-phosphorylating leak respiration (LEAK) was measured following additions of glutamate (G; 10 mM) and malate (M; 2 mM). Complex I-supported oxidative phosphorylation (OXPHOS) capacity was subsequently assessed by addition of ADP (D; 10 mM). Total OXPHOS capacity supported by complexes I and II was determined following addition of succinate (S; 20 mM).

Maximal electron transfer (ET) capacity was measured by stepwise titration of the uncoupler carbonyl cyanide m-chlorophenyl hydrazone (CCCP, U; 0.1 μM increments) until maximum O_2_ flux was achieved. Complex II-supported ET capacity was then determined following inhibition of complex I with rotenone (Rot; 0.5 μM). Finally, to determine experimental O_2_ baseline flux, residual oxygen consumption (ROX) was measured following inhibition of complex II with antimycin A (Ama; 2.5 μM) To prevent oxygen limitation, supplemental O_2_ was added throughout the assay to maintain dissolved O_2_ concentration above 250 μM.

Mitochondrial outer membrane integrity was assessed by addition of cytochrome c (10 µM), with preparations exhibiting >10% stimulation of O_2_ flux excluded from analysis. Air calibration of the polarographic O_2_ sensors was performed immediately prior to each SUIT protocol, and zero-O_2_ calibrations were performed on non-experimental days by dithionite addition. O_2_ flux values were corrected for instrumental background, O_2_ diffusion, sensor O_2_ consumption, and ROX to isolate mitochondrial O_2_ flux.

### Electrophysiology Recordings

SD was induced using elevated-K^+^ aCSF, prepared by replacing NaCl with KCl to achieve a final K^+^ concentration of 26 mM (Andrew et al. 2017) while maintaining physiological osmolarity. This modification was applied to each experimental condition (glucose, BHB, and sucrose).

Recordings were performed using an interface system at 34–37°C. Brain slices were continuously perfused (3.5 mL/min) with their respective carbogenated aCSF using a peristaltic perfusion system (PPS2 Multi Channel Systems). Following a 10-minute equilibration period, elevated-K^+^ (26 mM) aCSF was perfused for a total of 7 minutes or until SD occurred. Physiological-K^+^ aCSF (3.5 mM) was then restored for 10 minutes before repeating the SD induction protocol one additional time.

### Intrinsic Optical Signal Recording and Analysis

SD events were monitored using intrinsic optical signal (IOS) imaging. IOS images were acquired using a Lumenera INFINITY8-2M microscope camera (Teledyne Vision Solutions, Thousand Oaks, CA) mounted on an AmScope SM-6 Series zoom trinocular stereo microscope (AmScope, Irvine, CA). Camera exposure and gain settings were held constant throughout each recording.

Slices were illuminated using a broad-field battery-powered LED light source (Harbor Freight, Calabasas, CA). Images were acquired at 1 Hz with a resolution of 1608 × 1104 pixels using INFINITY ANALYZE 7 software and stored as uncompressed TIFF sequences. Spatial calibration was performed for each slice using a reference standard of known dimensions.

Image sequences were analyzed using a custom Python-based (Python Software Foundation, Beaverton, OR) analysis pipeline developed for SD identification and metric quantification. SD events were manually defined as start-to-end frame ranges and opened in a dedicated interface for preprocessing, segmentation, mask correction, ROI definition, spatial calibration, and metric export.

For each SD event, frames were preprocessed by applying Gaussian smoothing (σ = 0.5 pixels), subtracting a pixel-wise median baseline computed from the 20–30 frames preceding the SD event, and normalizing intensities to the global 1st–99th percentile range to enhance contrast of the SD-associated IOS. The neocortex was manually delineated as the region of interest (ROI). SD segmentation was performed using the SAM2.1 video predictor with the Hiera Base+ checkpoint (sam2.1_hiera_base_plus; Meta AI). Initial masks were generated from user-provided point prompts and brush annotations, then propagated through the selected event frame range. Propagated masks were manually reviewed and corrected before analysis.

Quantitative metrics were derived from the final binary masks. For each frame, the primary SD region was identified as the largest external contour and smoothed by retaining the 25 lowest Fourier harmonics. Propagation speed was calculated from frame-to-frame outward displacement of the SD boundary and converted to µm/s using the spatial calibration factor and image acquisition rate. Outward displacement values were thresholded to exclude small movements below 2 pixels, and the remaining displacements were averaged to obtain the mean boundary advance for the frame transition. When multiple disconnected SD regions were present, propagation speeds were calculated separately for each lineage-aware object and combined as an area-weighted mean. Frames with missing or invalid masks were excluded from speed calculations.

To account for slow reflectance changes associated with solution exchange during high-K^+^ treatment and washout, SD recovery analyses were performed on baseline-corrected IOS signals. For each event, relative reflectance change was calculated as ΔR/R_0_ = (R(t) − R_0_) / R_0_, where R_0_ was defined as the mean reflectance of the 30 frames preceding K^+^ application (baseline reflectance). A recruited-region trace was calculated from the cumulative SD mask within the neocortical ROI, while a reference trace was calculated from non-recruited ROI pixels. A piecewise baseline model representing the solution-associated reflectance component was generated from the reference trace and aligned to the recruited-region trace over the interval spanning SD onset to high-K^+^ washout. The SD-associated IOS signal was then isolated by subtracting this baseline model from the recruited-region trace.

Peak SD-associated IOS amplitude was defined as the largest signed residual deflection within 100 seconds of SD onset, located to sub-frame resolution by parabolic interpolation. Recovery half-time (T_50_) was defined as the interval from peak amplitude to the first time the residual signal returned to 50% of peak amplitude relative to the corrected baseline, assessed up to 250 seconds past the peak. Events were excluded from recovery analyses if no detectable SD signal was present (peak amplitude <0.5% ΔR/R_0_), if solution washout occurred before T_50_, or if technical issues compromised data quality.

### Statistical Analysis

Data are presented as mean ± standard error of the mean (SEM) or median and interquartile range, as appropriate. No differences were observed between male and female mice, and therefore data were presented as a single group. Parametric data assessed by Shapiro-Wilk testing were analyzed by unpaired two-tailed Welch’s *t* test or one-way ANOVA and nonparametric data by unpaired two-tailed Mann–Whitney *U* test or Kruskal–Wallis test. Contingency analyses were conducted by two-tailed Fisher’s exact test. All statistical analyses were performed using GraphPad Prism version 11.0 (Graph Pad Inc., San Diego, CA). IOS imaging data was analyzed using a custom-built Python graphical user interface as described above, as well as custom Python scripts. Sample sizes differed among metrics because progressively more stringent image-quality requirements were necessary for advanced IOS analyses, with propagation speed and residual ΔR/R_0_ measurements requiring higher signal quality and greater stability than SD occurrence or latency measurements. Investigators were not blinded to treatment assignment during data acquisition or analysis.

## Results

### BHB supports basal mitochondrial function without glucose

Following 4-hour maintenance of mouse neocortical tissue in either 10 mM BHB aCSF or 10 mM glucose aCSF, high-resolution respirometry was performed to assess mitochondrial respiratory function (Fig. 1). Across all respiratory states examined, tissue incubated with BHB exhibited significantly lower O_2_ flux (*J*_O2_) than tissue incubated with glucose (Fig. 1b-f). *J*_O2_ compensating for proton leak, proton slip, cation cycling and electron leak at high chemiosmotic potential (LEAK state), with recruitment of complex I when complex II and ATP synthase are inactive, was reduced in BHB-incubated tissue relative to glucose-incubated tissue (Fig. 1b).

**Fig. 1.**
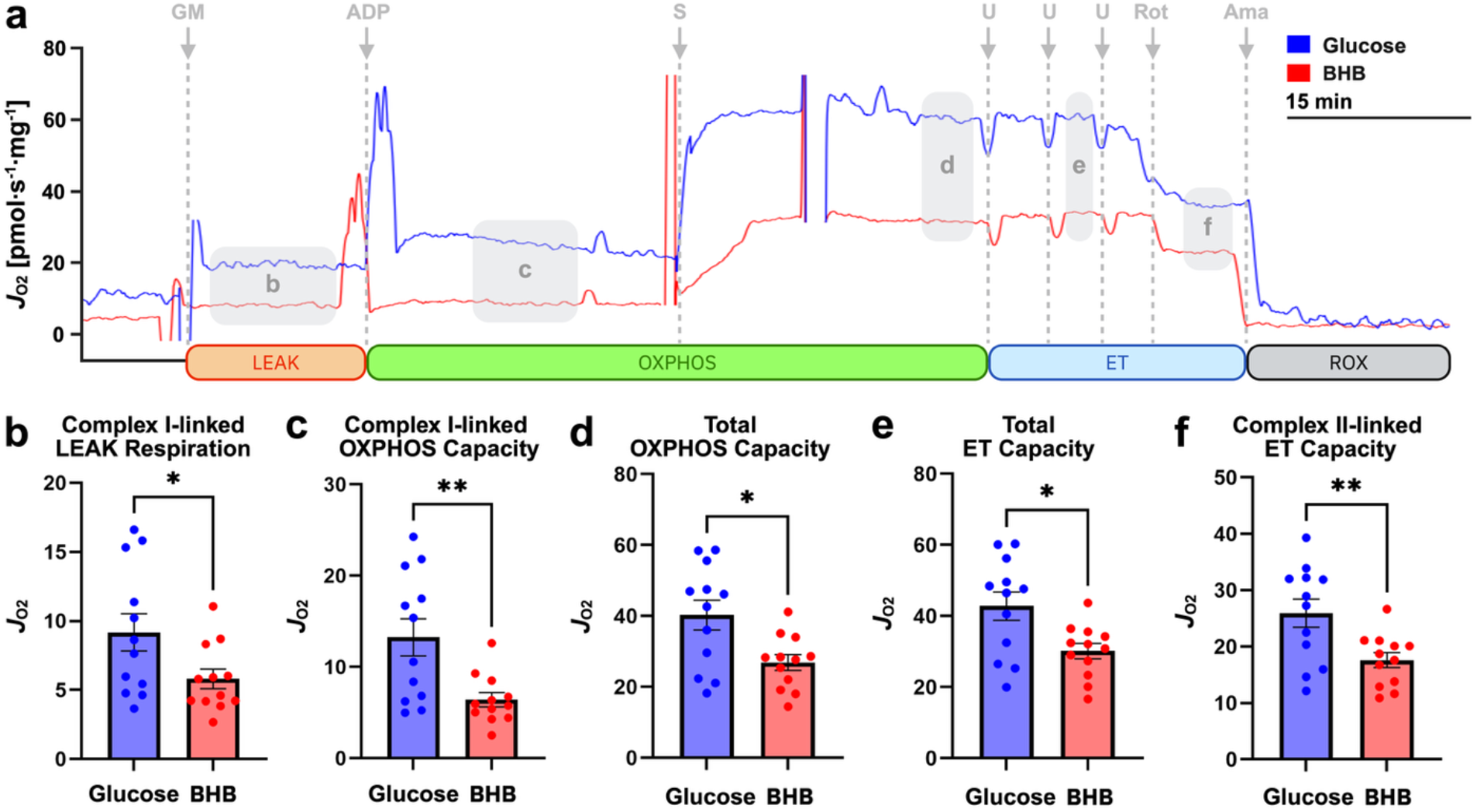
Mitochondrial oxygen flux in mouse neocortex is reduced following 4-hour incubation in 10 mM β-hydroxybutyrate (BHB) aCSF compared to 10 mM glucose aCSF. (a) Representative O_2_ flux (*J*_O2_) traces during a substrate-uncoupler-inhibitor-titration (SUIT) protocol depicting change in mitochondrial *J*_O2_ over time per unit tissue mass (pmol·s^-1^·mg^-1^). (b) Complex I-linked LEAK respiration, representing *J*_O2_ supported by electron entry through complex I in the absence of oxidative phosphorylation (OXPHOS) following addition of glutamate and malate (GM). (c) Complex I-linked OXPHOS capacity, representing maximal *J*_O2_ supported by electron entry through complex I coupled to ATP production following addition of ADP. (d) Total OXPHOS capacity, representing maximal *J*_O2_ from the additive contributions of complexes I and II electron supply pathways converging at the Q-junction coupled to ATP production following addition of succinate (S). (e) Total electron transfer (ET) capacity, the noncoupled state of maximum mitochondrial respiration following stepwise addition of the uncoupler CCCP (U). (f) Complex II-linked ET capacity, representing maximal *J*_O2_ supported by electron entry through complex II in the noncoupled state following addition of rotenone (Rot). Residual oxygen consumption (ROX), determined by Ama addition at the end of the SUIT protocol, was subtracted from *J*_O2_ values to isolate mitochondrial *J*_O2_ (baseline correction). Background corrected data are presented to remove the contribution of O_2_ backdiffusion and O_2_ consumption by the polarographic O_2_ sensor. Data are presented as mean ± SEM (unpaired two-tailed Welch’s *t* test, *n* = 12). \**p* < 0.05, \*\**p* < 0.01. Created in BioRender. Parrish, R. (2026) https://BioRender.com/kit9f5b.

With recruitment of ATP synthase in the ADP-activated state of oxidative phosphorylation (OXPHOS), *J*_O2_ remained lower following BHB incubation in both the complex I-supported OXPHOS state (Fig. 1c) and the complex I and II-supported OXPHOS state (Fig. 1d). Similarly, *J*_O2_ associated with maximal electron transfer capacity (ET capacity) in the uncoupled state was reduced following BHB incubation when supported by complexes I and II (Fig. 1e) and by complex II alone (Fig. 1f). While *J*_O2_ following BHB treatment was reduced compared to glucose controls, this data suggests that BHB can preserve mitochondrial function during extreme glucose deprivation.

### BHB alone can supply the energy required for highly energetically demanding brain events

Neocortical brain slices were subjected to a high-K^+^ bath to induce SDs (Fig. 2a). Slices incubated in BHB or glucose aCSF fully recovered from SD within 10 minutes and experienced an additional SD upon repeat high-K^+^ stimulation (Fig. 2b). Additional high-K^+^ stimulation was unable to elicit SDs in fuel-deprived slices (Fig. 2bii).

**Fig. 2.**
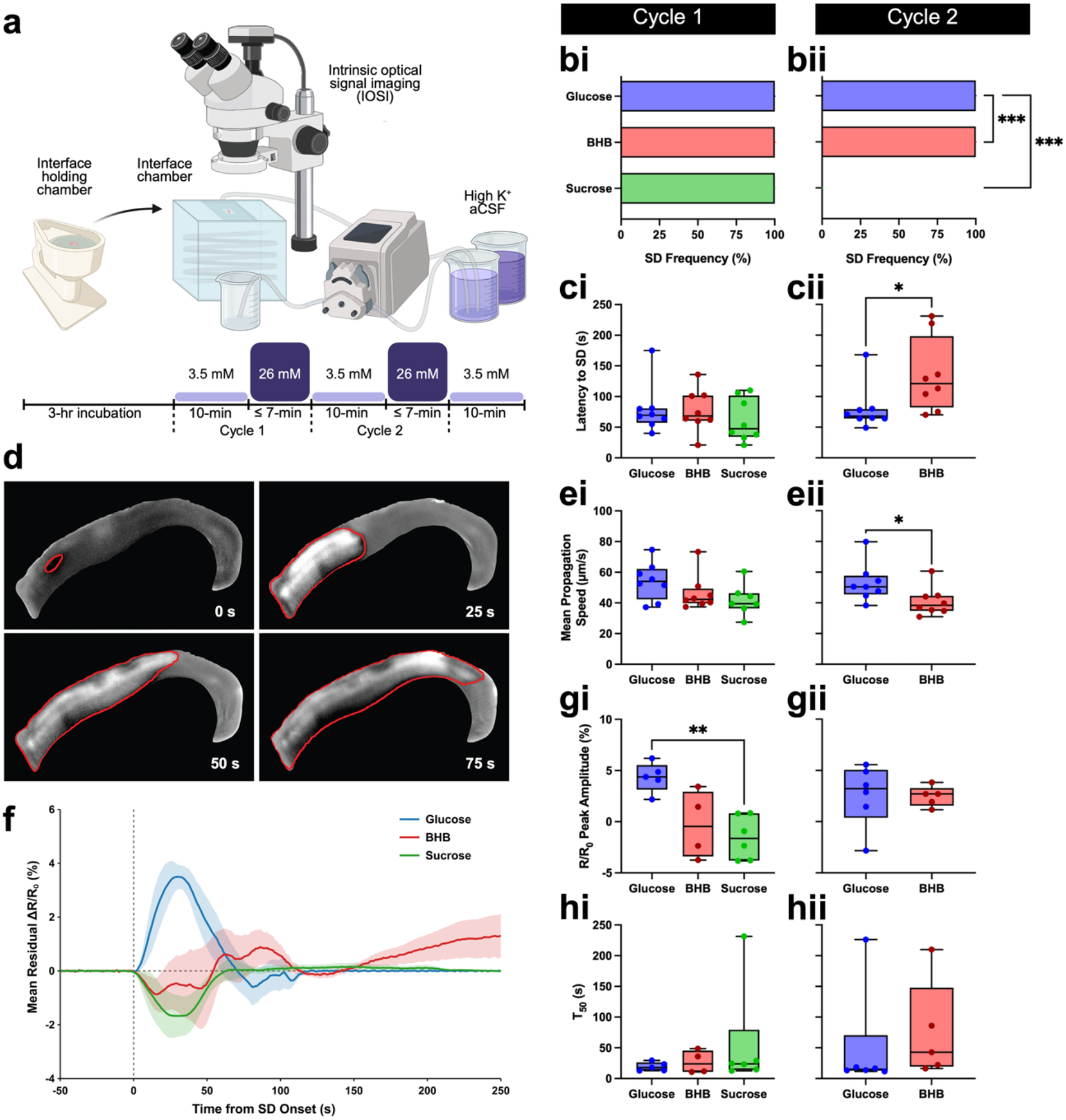
Neocortical brain slices supplied with BHB in the absence of glucose can undergo and recover from energetically demanding spreading depolarizations (SD). (a) Experimental paradigm for sequential SD induction. (b) SD occurrence rate upon high-K^+^ stimulation in media containing 10 mM glucose, 10 mM BHB, or 10 mM sucrose (fuel-deprivation) in (bi) cycle 1 and (bii) cycle 2 (two-tailed Fisher’s exact test, *p* < 0.001, *n* = 8 per group). (c) Latency to SD after 26 mM high K^+^ insult for (ci) cycle 1 (Kruskal–Wallis test, *p* = 0.5628, *n* = 8 per group) and (cii) cycle 2 (two-tailed Mann–Whitney *U* test, *p* = 0.027, *n* = 8 per group). (d) Representative intrinsic optical signal (IOS) images of SD propagating across the neocortex displayed at 25-second intervals. Red lines indicate the SD border. (e) Mean propagation speed of each SD event for (ei) cycle 1 (Kruskal–Wallis test, *p* = 0.1842 *n* = 8 for glucose, *n* = 8 for BHB, *n =* 7 for sucrose) and (eii) cycle 2 (two-tailed Welch’s *t* test, *p* = 0.0448, *n* = 8 per group). (f) Mean residual change in reflectance (ΔR/R_0_) for the area recruited by SD over time for cycle 1 (*n* = 5 for glucose, *n* = 4 for BHB, *n =* 6 for sucrose). The solid line represents the mean for each group, while the shaded region displays the ± standard error of the mean (SEM). (g) Peak amplitude of SD-associated IOS relative to baseline reflectance prior to SD (ΔR/R_0_) for (gi) cycle 1 (one-way ANOVA with Tukey’s post-hoc tests, *p* = 0.0035, *n* = 5 for glucose, *n* = 4 for BHB, *n* = 6 for sucrose) and (gii) cycle 2 (two-tailed Welch’s *t* test, *p* = 0.9299, *n* = 6 for glucose, *n* = 5 for BHB). (h) Time from peak amplitude to return to 50% of the signed peak amplitude relative to baseline following SD for (hi) cycle 1 (Kruskal–Wallis test, *p* = 0.8597, *n* = 5 for glucose, *n* = 4 for BHB, *n* = 6 for sucrose) and (hii) cycle 2 (two-tailed Mann–Whitney *U* test, *p* = 0.1775, *n* = 6 for glucose, *n* = 5 for BHB). Boxplots display the median and interquartile range, with whiskers extending to the minimum and maximum values. \**p* < 0.05, \*\**p* < 0.01, \*\*\**p <* 0.001. Created in BioRender. Parrish, R. (2026) https://BioRender.com/1feya74.

Within the first elevated-K^+^ cycle, latency to SD and SD propagation speed did not vary significantly between groups (Fig. 2ci,ei). However, following a second SD induction, BHB-treated slices showed a significantly longer latency and slower rate of SD propagation than glucose-treated slices (Fig. 2cii,eii).

IOS analysis further revealed that all treatment groups returned to baseline reflectance following the first SD (Fig. 2f). There was also no difference in ΔR/R_0_ peak amplitude between glucose- and BHB-incubated slices (Fig. 2g). However, slices incubated in glucose aCSF but not BHB aCSF, demonstrated significantly higher ΔR/R_0_ peak amplitude than fuel-deprived slices (Fig. 2gi). Time to return to 50% baseline reflectance did not differ significantly between any groups (Fig. 2h). Collectively, this data suggests that BHB can support extreme brain tissue demands without glucose.

## Discussion

Glucose has classically been viewed as a requisite energetic substrate for neural tissue. Neurons, however, demonstrate metabolic flexibility in their capacity to utilize a variety of substrates, including lactate and ketone bodies (Dienel 2019). The present study examined whether BHB can independently support brain tissue metabolism in the absence of exogenous glucose. We found that acute reliance on BHB reduced mitochondrial *J*_O2_ across all respiratory states measured by high-resolution respirometry relative to tissue subsisting primarily on glucose. Despite this reduction, BHB-supported tissue demonstrated the ability to generate and recover from repeated SDs, metabolically demanding events that require rapid restoration of transmembrane ion gradients. Furthermore, repeated SD induction revealed a longer latency to SD onset and slower propagation in BHB-treated tissue relative to glucose-treated tissue. Together, these findings suggests that BHB alone can sustain brain metabolic function for several hours and may reduce tissue susceptibility to repeated SD.

A notable finding was the reduction in mitochondrial *J*_O2_ associated with mitochondrial leak respiration, oxidative phosphorylation capacity, and electron transfer capacity following incubation in BHB compared with glucose. Different catabolic substrates often exhibit different rates of respiration supporting similar power outputs due to adaptive differences in coupling ef,iciency, proton leak, ATP/O_2_ stoichiometry, and the thermodynamic state of ATP hydrolysis (ΔG_ATP_) (Gnaiger 2020; Schmidt et al. 2021). Indeed, ketone body metabolism has been proposed to alter mitochondria bioenergetics through increasing mitochondrial coupling efficiency and ATP flux (Holstein et al. 2025), increasing ΔG_ATP_ (Veech 2004; Veech et al. 2001), reducing free radical damage (Achanta and Rae 2017; Jarrett et al. 2008; Veech et al. 2001), and improving bioenergetic resilience during periods of heightened energetic demand (Llorente-Folch et al. 2023). Thus, the reduction in *J*_O2_ in mitochondria acutely adapted to BHB usage may reflect a more efficient metabolic strategy (Saito et al. 2022; Veech et al. 2001) compared to glucose rather than a reduction in ATP generative capacity.

The observation that BHB-supported tissue remained capable of generating and recovering from repeated SDs upon high-K^+^ stimulation supports the conclusion of metabolic sufficiency. SD involves a massive redistribution of ions and neuroactive compounds across neuronal and glial membranes. Restoration of ionic homeostasis requires substantial ATP consumption, primarily through the activation of Na^+^/K^+^ ATPases and other ion transport systems (Lindquist 2024; Ricks et al. 2025). Because recovery from SD is one of the most significant energetic challenges encountered by brain tissue, the ability of BHB-treated slices to undergo a second SD following recovery from the first indicates that ketone-supported tissue retained sufficient metabolic capacity to restore ionic homeostasis.

Initial latency from high-K^+^ insult to SD induction and average propagation speed of the SD wavefront were similar between glucose- and BHB-treated slices. Interestingly, for repeated SDs, BHB-treated tissue exhibited prolonged latency to SD onset and slower average SD propagation. Propagation of the SD depends on the ability of the SD wavefront to recruit adjacent neurons into the regenerative wave. Consequently, both increased latency and reduced propagation velocity are generally interpreted as markers of reduced SD susceptibility (Aiba and Noebels 2021; Dreier 2011; Jansen et al. 2024; Vitale et al. 2025). Energy deprivation is a common SD induction paradigm (Ricks et al. 2025); therefore, BHB’s therapeutic effect may be brought about by increased energy availability during these extreme brain perturbations.

Analysis of SD-associated IOS further support the conclusion that BHB can sustain neural tissue function in the absence of exogenous glucose. Following the initial SD, IOS returned to baseline in all brain slices. Neither peak IOS amplitude nor time to 50% recovery differed between glucose- and BHB-treated slices, suggesting that ketone-supported tissue can restore ionic homeostasis and reverse SD-associated cellular swelling at a rate comparable to glucose-supported tissue. Importantly, despite similar recovery kinetics following the first SD, sucrose-treated brain tissue was unable to support a subsequent SD, whereas slices incubated with glucose or BHB could generate and recover from repeated SD events.

Together, these data suggest β-hydroxybutyrate is sufficient to support essential brain tissue metabolism and function in the absence of exogenous glucose, demonstrating cerebral metabolic flexibility.

## Statements & Declarations

### Funding

This research was funded by internal support provided by Brigham Young University.

### Competing Interests

Benjamin T. Bikman is an advisor for Unicity International and receives royalties for his role with them, along with royalties from the sale of a book about insulin resistance. Benjamin T. Bikman, Paul R. Reynolds, and R. Ryley Parrish are advisors for Ketone Labs. All authors declare there are no additional conflicts of interest.

### Author Contributions

All authors contributed to the study conception and design. Material preparation, data collection, and analyses were performed by Rachel Ricks, Dallin S. Nevers, Tyler Poulos, T. Luke Shafer, and Clay Dunford. IOS imaging analysis software was designed by Clay Dunford. The first draft of the manuscript was written by Rachel Ricks, Dallin S. Nevers, Tyler Poulos, T. Luke Shafer, Clay Dunford, and R. Ryley Parrish, and all authors commented on previous versions of the manuscript. All authors read and approved the final manuscript.

### Data Availability

The raw datasets generated during the current study are not publicly available due to lack of standard repository for this type of data but are available from the corresponding author upon reasonable request.

### Ethics Approval

All animal experiments were conducted in accordance with the National Institutes of Health and the Brigham Young University Institutional Animal Care and Use Committee.

## Notes

### Competing Interest Statement

BTB is an advisor for Unicity International and receives royalties for his role with them, along with royalties from the sale of a book about insulin resistance. BTB, PRR, and RRP are advisors for Ketone Labs.

### Summary of Updates

Figure 2 revised to correct formatting error.

